# Human TBC1 domain-containing kinase is a class I multidomain pseudokinase

**DOI:** 10.64898/2026.04.02.716191

**Authors:** Shiwangi Maurya, Lilly E. Cheek, Anthony T. Iavarone, Wen Zhu

## Abstract

TBCK-related encephalopathy (TBCKE) is a neurodevelopmental disorder associated with biallelic mutations in *TBCK*. Despite the increasing number of reported cases worldwide, the biochemical and biophysical properties of TBCK remain unclear, hindering molecular understanding of its role in disease. Here, we present the successful expression, purification, and biochemical characterization of full-length human TBCK produced in *Spodoptera frugiperda* cells. Biochemical and biophysical analyses reveal that the catalytically inactive pseudokinase domain of TBCK lacks nucleotide binding, consistent with the absence of the canonical VAIK, HRD, and DFG motifs required for catalysis. These findings support that TBCK is a class I pseudokinase and provide a foundation for future structural and functional studies to elucidate its biological role.

## 1. Introduction

TBCK encephaloneuronopathy (TBCKE), also known as TBCK syndrome, or infantile hypotonia with psychomotor retardation and characteristic facies type 3, is an autosomal recessive inborn neurodevelopmental disorder ^1^. Since its initial recognition as a distinct clinal entry in 2015, TBCKE has been reported in more than 60 individuals worldwide and is now recognized as a severe inborn error of neurodevelopment ^2^. Affected individuals present with profound global developmental delay and combined intellectual and motor impairment, commonly accompanied by macrocephaly, axial hypotonia, seizures, and marked deficits in speech and voluntary movement ^3–5^. Genetic analyses have established that TBCKE is caused by biallelic pathogenic variants in the gene encoding TBC1 domain–containing kinase (TBCK)^1,6–8^.

Recent studies have identified TBCK as an essential component of the FERRY complex, a multiprotein assembly that functions as a Rab5A effector in early endosomal trafficking ^9,10^. The FERRY complex has been proposed to mediate mRNA binding and transport along endosomal pathways, suggesting a direct molecular link between TBCK function, vesicular trafficking, and neuronal development. Despite these advances, the biochemical activity and biophysical properties of TBCK itself remain poorly defined, limiting mechanistic understanding of how disease-linked variants disrupt neurodevelopmental processes.

TBCK is predicted to comprise three distinct domains arranged from N- to C-terminus: a pseudokinase domain, a Tre2-Bub2-Cdc16 (TBC) domain, and a rhodanese-like domain, connected by short linker regions (**Figure 1a**). Bioinformatic analysis indicates that the pseudokinase domain lacks the canonical catalytic motifs required for serine/threonine kinase activity, including the glycine-rich loop, VAIK, HRD, and DFG motifs (**Figure 1b**), consistent with the classification of TBCK as a class I pseudokinase. Although sequence analysis is informative for identifying hallmark features of pseudokinases, numerous annotated pseudokinases have been shown to retain unexpected ligand-binding capabilities or residual enzymatic activity through the substitution of alternative residues within or adjacent to the canonical active site ^11,12^. Therefore, experimental validation is essential to establish the functional classification and biochemical properties of individual pseudokinases.

**Figure 1.**
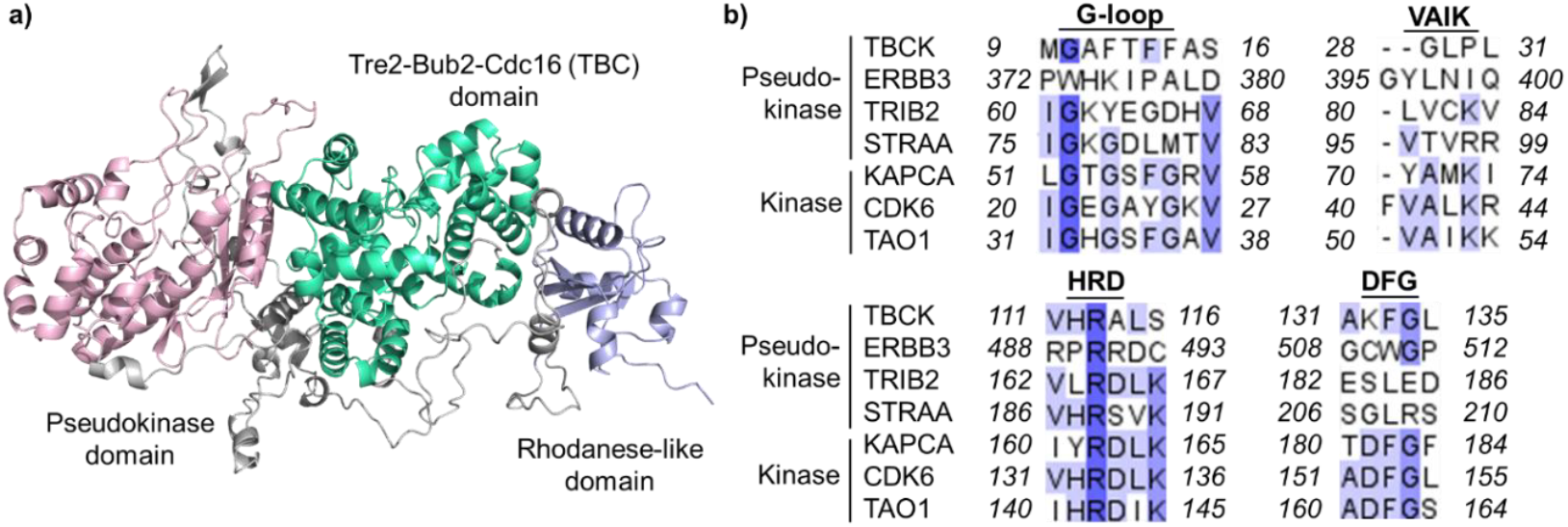
Structural and sequence alignment analysis of TBCK. **a)** Domain organization and predicted AlphaFold structure (Q8TEA7-F1) of full-length human TBCK. The model shows the N-terminal pseudokinase domain (light pink), central Tre2-Bub2-Cdc16 (TBC) domain (green), and C-terminal rhodanese-like domain (light purple), arranged from N- to C-terminus and connected by short linker regions (grey). **b)** Sequence alignment results of TBCK with other pseudokinases and active kinases showing missing residues in key motifs, including the G-loop, VAIK, HRD, and DFG motifs. The UniProt codes for these aligned sequences include representative pseudokinases: human TBCK (P21860), human ERBB3 (Q92519), human TRIB2 (Q92519), and human STRAA (Q7RTN6), as well as catalytically active protein kinases: human KAPCA (P17612), human CDK6 (Q00534), and human TAO1 (Q9H2KQ).

In this study, we report successful expression, purification, and *in vitro* biochemical characterization of TBCK. These foundational analyses provide a framework for directly testing the functional potential of the TBCK pseudokinase domain and establish a biochemical basis for understanding how TBCK dysfunction contributes to its biological role in cells, with potential implications for future diagnostic and therapeutic strategies for TBCKE.

## 2. Materials and Methods

### 2.1 Construct design and bacmid preparation

The canonical amino acid sequence of human TBCK was obtained from UniProt (Q8TEA7). A TEV-cleavable His-10 tag (AGENLYFQGHHHHHHHHHH) was added to the C-terminus of the TBCK sequence. The DNA sequence of TBCK was codon-optimized for heterologous expression in *Spodoptera frugiperda (*Sf9) insect cells using a baculovirus expression system. The *TBCK* gene was synthesized and cloned into the pFastBac1 vector between the BssHII and XhoI restriction sites by GenScript (Piscataway, NJ). The sequence of the inserted gene was confirmed by Sanger sequencing at the Florida State University DNA Sequencing Facility. The pFastBac-TBCK plasmid was transformed into DH10Bac competent cells and plated onto LB agar plates containing 100 mM isopropyl β-D-1-thiogalactopyranoside (IPTG), 20 mg/mL 5-bromo-4-chloro-3-indolyl β-D-galactopyranoside (X-gal), 7 μg/mL gentamicin, 10 μg/mL tetracycline, and 50 μg/mL kanamycin, and incubated for 48 hours at 37 °C. Six white colonies were selected, streaked onto a fresh LB agar plate containing the above-mentioned reagents, and incubated for another 48 hours at 37 °C. After overnight incubation, a single colony was inoculated into 5 mL LB media supplemented with 50 μg/mL kanamycin, 7 μg/mL gentamicin, and 10 μg/mL tetracycline and incubated overnight at 37 °C, 250 rpm. The bacmid was isolated from the overnight cell culture using a method adapted from the previously published protocol^13^. The purified bacmid DNA was stored at -20 °C before transfection into Sf9 cells.

### 2.2 Baculovirus production and amplification

Sf9 insect cells were seeded at a density of 1.5 × 10^6^ cells per well in a 6-well plate containing ESF 921 insect cell culture medium supplemented with 1% fetal bovine serum (FBS) and incubated for 1 hour at 28 °C to allow cell attachment. For transfection, 2 μg of purified recombinant bacmid DNA was mixed with polyethylenimine (PEI: DNA ratio, 1:4) in 300 μL of 150 mM NaCl. This mixture was incubated at room temperature for 30 minutes. The resulting transfection mixture was then added dropwise to the cells to ensure uniform distribution, and the cells were incubated at 28 °C. After 6 days, the culture supernatant containing the P0 viral stock was collected by centrifugation at 500 × g for 5 minutes, filtered through a 0.22 μm syringe filter, and stored at 4 °C for subsequent amplification to produce the P1 virus stock. For P1 virus amplification, 30 mL of Sf9 cells at a density of 2 × 10^6^ cells/mL were infected with 1 mL of P0 virus, and cell density and viability were monitored daily. After approximately 90 hours, when cell viability dropped to ∼57%, the P1 virus was harvested by centrifugation at 500 × g for 5 minutes and filtered through a 0.22 μm syringe filter, supplemented with 1% FBS, and stored at 4 °C for subsequent protein expression.

### 2.3 Expression of TBCK

Sf9 cells at a density of 2 × 10^6^ cells/mL were infected with the P1 viral stock at a multiplicity of infection (MOI) of 1. The MOI was determined by plaque assay^14^. The culture was maintained in suspension by shaking at 120 rpm at 28 °C for 60 hours. After incubation, cells were harvested by centrifugation at 500 × g for 10 minutes, and the pellet was stored at -80 °C until purification.

### 2.4 Purification of TBCK

All the purification steps were performed at 4 °C. The cell pellet collected from the expression culture was resuspended in Buffer A (50 mM EPPS, 150 mM NaCl, 25 mM imidazole, 5 mM TCEP, pH 8) on ice and sonicated for 3 minutes (1 second ON/1 second OFF, 25% amplitude). The lysate was centrifuged at 12,000 × g for 45 minutes and filtered through a 0.45 μm syringe filter. The filtered lysate was loaded on a pre-equilibrated Ni-NTA column with Buffer A. The column was washed with 30 mL of Buffer B (50 mM EPPS, 150 mM NaCl, 60 mM imidazole, 5 mM TCEP, pH 8), and protein was eluted with Buffer C (50 mM EPPS, 150 mM NaCl, 250 mM imidazole, 5 mM TCEP, pH 8).

Fractions were pooled, concentrated using an Amicon Ultra 30K centrifugal filter, and buffer-exchanged into Buffer D (50 mM EPPS, 150 mM NaCl, 10% glycerol, 5 mM TCEP, pH 8) using a PD-10 desalting column. The protein was further purified by size-exclusion chromatography (SEC) on a HiPrep 16/60 Sephacryl S-100 HR column equilibrated with filtered, degassed Buffer E (50 mM EPPS, 150 mM NaCl, 5 mM TCEP, pH 8.0). Pure TBCK fractions confirmed by SDS-PAGE were pooled, concentrated, buffer exchanged into Buffer D, flash-frozen in liquid nitrogen, and stored at -80 °C.

### 2.5 Protein concentration determination

UV-Vis spectra were recorded using an Agilent Cary 60 UV-Vis spectrophotometer. Protein samples were diluted in Buffer F (20 mM potassium phosphate, 20 mM NaCl, 2 mM DTT, pH 8.0). Amino acid analysis was performed at the Molecular Structure Facility, University of California, Davis. Samples were first eluted twice with 200 μL of 5:50 (v/v) formic acid/acetonitrile, and the two eluates were combined. The pooled fractions were transferred to a hydrolysis vial and evaporated to dryness. The dried residue was resuspended in 200 μL of 6 N HCl containing 1% (w/v) phenol, and hydrolysis was carried out at 110 °C for 24 hours. After cooling to room temperature, the samples were dried again. The dried samples were then reconstituted in Norleucine (NorLeu) dilution buffer to a final volume of 2.0 mL. For amino acid analysis, 50 μL of the reconstituted hydrolysate was injected into a Hitachi LA8080 ion-exchange amino acid analyzer equipped with post-column ninhydrin detection using NorLeu as the internal standard for quantification. Routine protein concentrations were determined by Bradford assay and corrected using a scaling factor of 0.75 derived from amino acid analysis.

### 2.6 Analytical size-exclusion chromatography (SEC)

A Superdex 200 Increase 5/150 GL column was used to determine TBCK oligomerization state in solution. TBCK was injected onto the SEC column equilibrated with Buffer E, at a flow rate of 0.22 mL/min. Sigma Protein Standard Mix (15-600 kDa) was run under identical conditions to generate a standard curve. The molecular weight of TBCK was estimated from its elution volume relative to the standard curve.

### 2.7 Mass spectrometry (MS) analysis of TBCK

9 μM TBCK was equilibrated in Buffer E before being partially denatured in Buffer G (3 M guanidine hydrochloride, 1 M citric acid, 3 mM TCEP, pH 1). It was then subjected to proteolysis with 2 mg/mL pepsin for 10 minutes on ice. After digestion, the samples were flash-frozen in liquid nitrogen and stored until liquid chromatography-tandem mass spectrometry (LC-MS/MS) analysis, performed as described previously^15^. Briefly, samples were analyzed using a 1200 Series LC system (Agilent Technologies, Santa Clara, CA) that was connected in line with an LTQ-Orbitrap-XL mass spectrometer equipped with an electrospray ionization source (Thermo Fisher Scientific, Waltham, MA). The LC column compartment was equipped with an Acquity UPLC BEH C8 column (length: 100 mm, inner diameter: 1.0 mm, particle size: 1.7 μm, stock-keeping unit: 186002876, Waters, Milford, MA). Data acquisition and analysis were performed using Xcalibur (version 2.0.7) and Proteome Discoverer (version 1.3, Thermo) software. The sequence coverage of TBCK was visualized using HDX-WorkBench software ^16^.

### 2.8 Circular dichroism (CD) measurements

Purified TBCK was buffer exchanged into Buffer F and diluted to 0.25 mg/mL. CD spectra were recorded at 20 °C using a Chirascan VX spectrophotometer equipped with a temperature controller and a 0.1 cm path-length quartz cuvette. Spectra were collected from 190-280 nm (1 nm bandwidth, 25 nm/min scan speed, 0.2 nm data pitch). Buffer spectra were subtracted. The measurements were performed in triplicate. Secondary structure analysis was performed using BeStSel ^17^. For thermal denaturation, the CD signal at 208 nm was monitored as the temperature increased from 20 to 90 °C at 1 °C/min. The melting curve was fitted to a sigmoidal Boltzmann function to determine the apparent melting temperature (T_m_).

### 2.9 Differential Scanning Fluorimetry (DSF) assay

Thermal stability of TBCK was assessed in the absence and presence of nucleotides using the GloMelt assay (Biotium, Fremont, CA). Reactions containing 0.1 mg/mL TBCK, GloMelt dye, and 50 mM EPPS, 150 mM NaCl, pH 8.0. For the MgCl_2_ binding assay, 10 mM MgCl_2_ was used with apo-TBCK. For the nucleotide binding assay, ATP or GTP was titrated into TBCK at 1 mM, 5 mM, and 10 mM in the presence of 10 mM MgCl_2_. ADP and GDP were titrated into the TBCK solution in the presence of 10 mM potassium phosphate and 10 mM MgCl_2_. Reactions (20 μL) were transferred to a 96-well plate, incubated on ice for ∼1 hour, and thermal unfolding was monitored on a QuantStudio 7 Flex Real-Time PCR system (SYBR Green channel). The temperature was ramped from 20 to 95 °C at 0.05 °C/s. Data from biological duplicates were analyzed by plotting the first derivative of fluorescence (dF/dT) *vs*. temperature. T_m_ was determined as the peak maximum. Ligand-included ΔT_m_ was calculated by comparing melting temperatures of TBCK in the absence or presence of nucleotide conditions relative to TBCK-only controls. All measurements were performed in triplicate.

### 2.10 ATPase activity assay

The ability of the pseudokinase domain to hydrolyze the gamma phosphate group of ATP or GTP was assessed using a continuous, coupled activity assay adapted from the EnzChek Pyrophosphate Assay Kit (Molecular Probes, Eugene, OR). A reaction mixture was prepared in Assay Buffer (100 mM EPPS, 150 mM NaCl, 2 mM DTT, pH 8) containing 5 mM ATP or 5 mM GTP, 10 mM MgCl_2_, 0.2 mM 2-amino-6-mercapto-7-methylpurine ribonucleoside, and 1.5 U/mL purine nucleoside phosphorylase. Upon the addition of 30 nM TBCK, the absorbance at 360 nm was measured every 0.02 seconds for 6 minutes at 37 °C to assess potential phosphate production. Negative controls in which ATP or GTP was omitted from the reaction mixture were prepared and measured in the same manner. Human asparagine synthetase (50 nM) was used as a positive control because its ATP-hydrolyzing activity is well-defined under similar assay conditions using these reagents ^18,19^. In this positive control, in addition to the reaction mixture as described previously, 12.5 mM L-aspartate, 100 mM ammonium chloride, and 0.09 U/mL inorganic pyrophosphatase were also added to the mixture. All assays were performed in triplicate and visualized using GraphPad Prism (GraphPad Holdings, La Jolla, CA).

## 3. Results

### 3.1 Expression and purification of TBCK

Approximately 1 g of wet cell pellet was obtained from a 50-mL expression culture, yielding ∼2 mg purified TBCK protein. SDS-PAGE (**Figure 2a**) showed that TBCK migrated as a single band, and size-exclusion chromatography (SEC) indicated an apparent molecular weight of 103 kDa, consistent with a predominantly monomeric species in solution (**Figure 2b**). The UV-Vis spectrum of purified TBCK showed a characteristic 280 nm peak, and no other absorption features were observed (**Figure 2c**). The purified TBCK protein was subjected to limited proteolysis with pepsin, followed by LC-MS/MS analysis. The resulting peptide library comprised 480 unique peptides (**Figure 2d**), achieving 78.2% sequence coverage. Mapping these peptides onto the human TBCK sequence revealed two regions with reduced peptide coverage, residues 227-237 and 660-680, corresponding to the pseudokinase and TBC domains, respectively (**Figure 2e**).

**Figure 2.**
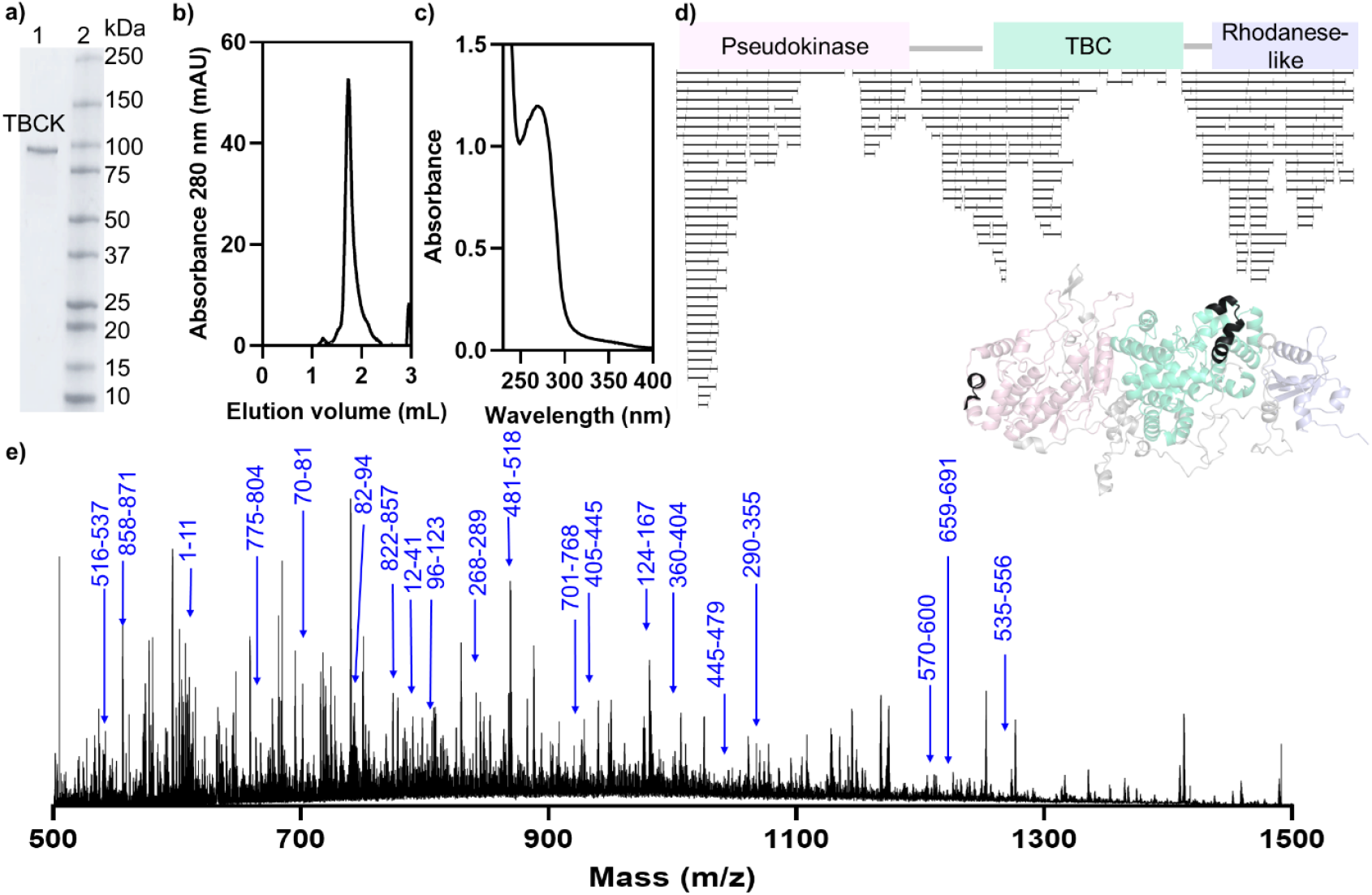
a) SDS-PAGE analysis of purified TBCK. Lane 1: Purified TBCK, Lane 2: Protein marker. b) Analytical size exclusion chromatography (SEC) profile. c) UV-Vis spectrum of purified TBCK showing a characteristic peak at 280 nm. d) Peptide mapping of limited proteolysis. Top, representative peptides are shown in black bars. Undetectable peptides are located in pseudokinase (227-237) and TBC (660-680) domains. Bottom, segments 227-237 and 660-680 are highlighted in black on the Alphafold-predicted structure to show the undetectable regions after proteolysis. e) Mass spectrometry analysis of TBCK after limited proteolysis with pepsin. Representative peptides are highlighted in blue.

### 3.2 TBCK functions as a pseudokinase lacking nucleotide-binding activity

To understand the secondary structure of TBCK, we performed the CD measurements at various temperatures (**Figure 3a**). The BestSel analysis of the spectrum at 20 °C assigned a predominantly α-helical secondary structure, with 64.8% helix, 9.4% antiparallel β-sheet, and 25.7% similar to the pseudokinase and TBC domain folds predicted by AlphaFold 4 ^20,21^ (**Figure 3b**). The temperature-dependent CD profile indicates a sigmoidal thermal unfolding transition with melting temperature (T_m_) of 47.31 ± 0.02 °C **(Figure 3c)**.

**Figure 3.**
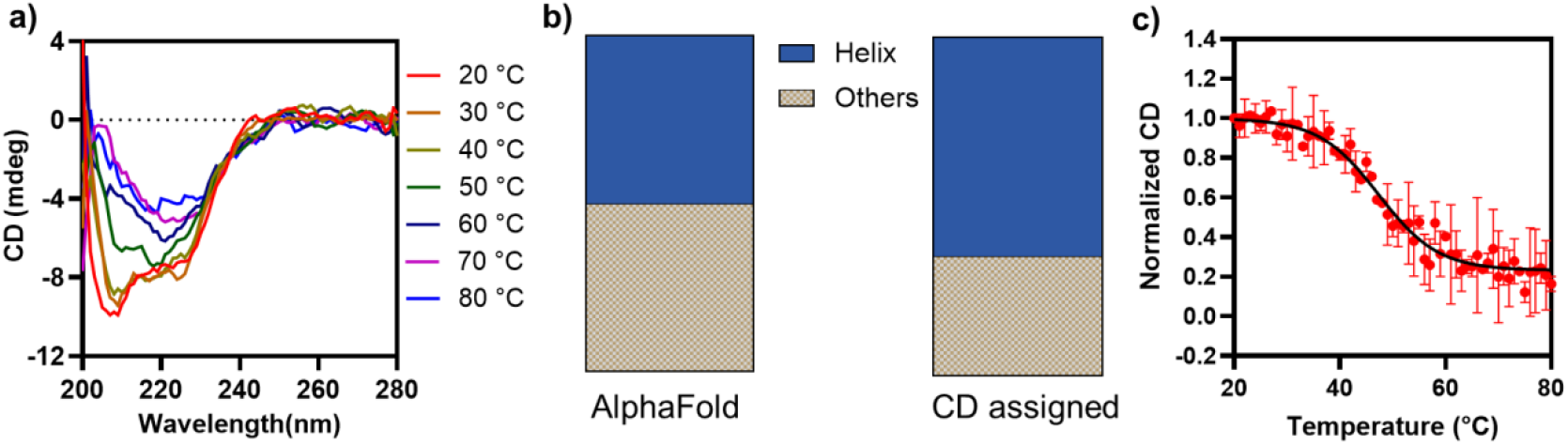
Circular dichroism (CD) profile of TBCK. a) Temperature-dependent CD profile. b) AlphaFold-predicted secondary structures and the BestSel secondary structure analysis calculated from the CD trace at 20 °C. c) Melting temperature determined by CD.

Class I pseudokinases lack the conserved motifs required for nucleotide and divalent cation binding ^22^. To test whether TBCK is a class I pseudokinase, we assessed nucleotide- and divalent cation-binding using DSF, as ligand binding is expected to increase protein thermal stability at elevated temperature. The T_m_ of apo TBCK determined by DSF is 47.3 ± 0.6 °C, consistent with the results obtained from temperature-dependent CD spectroscopy. In the presence of MgCl_2_, TBCK showed no change in the T_m_ (47.2 ± 0.6 °C) (**Figure 4a**). In the presence of ATP, the T_m_ of apo TBCK was recorded as 47.2 ± 0.5 °C, 47.5 ± 0.0 °C, and 47.5 ± 0.4 °C across nucleotide concentrations of 1, 5, and 10 mM, respectively, indicating no ATP-dependent stabilization (**Figure 4b**). GTP produced a slight decrease in T_m_ at 1 mM (45.4 ± 0.5 °C), with values of 46.8 ± 0.8 °C and 47.3 ± 0.2 °C at 5 and 10 mM, respectively, compared to apo TBCK (**Figure 4c**). The presence of ADP and phosphate (Pi) resulted in marginal increases in T_m_ (47 ± 1 °C, 48.5 ± 0.2 °C, 48.5 ± 0.2 °C) across the tested concentrations (**Figure 4d**). Similarly, replacing ADP by GDP showed minor effects, yielding T_m_ values of 46.5 ± 0.5 °C, 47.1 ± 0.3 °C, and 47.3 ± 0.9 °C at 1, 5, and 10 mM, respectively (**Figure 4e**). All observed T_m_ changes compared to the apo TBCK are approximately 1 °C and fall within the experimental error. These results, therefore, indicate that none of the tested nucleotides measurably stabilizes or destabilizes the global thermal stability of TBCK under the conditions examined, consistent with TBCK functioning as a class I pseudokinase.

**Figure 4.**
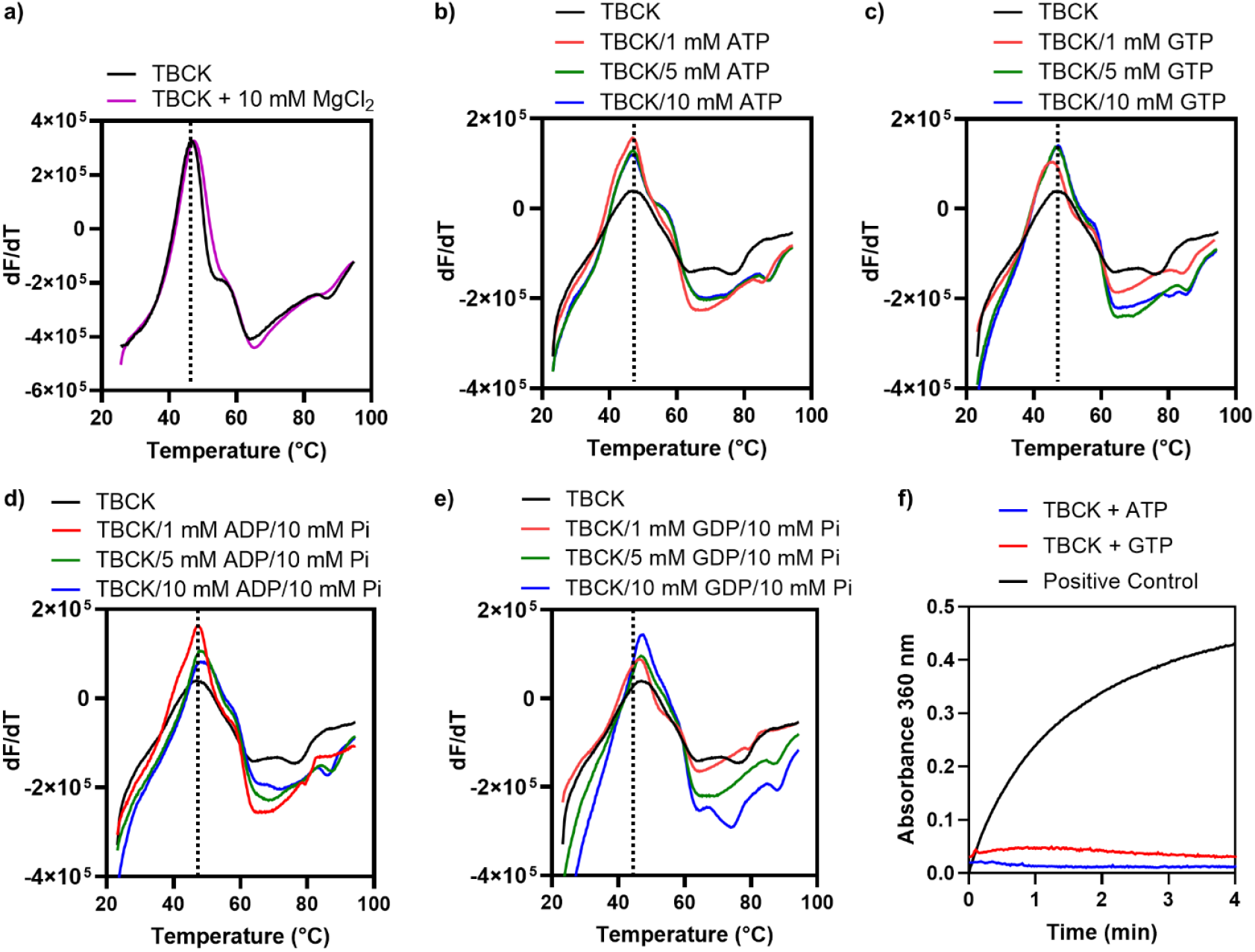
Probing the ligand binding ability of TBCK using DSF and activity assays. For DSF assays, first derivative plots (dF/dT *vs*. temperature) of the thermal unfolding of TBCK in the presence of 1 mM (red), 5 mM (green), and 10 mM (blue) nucleotides were plotted in comparison with apo TBCK. Each curve shown here is an average of biological duplicates. Solid black traces represent apo TBCK. Thermal unfolding is shown in the presence of a) 10 mM MgCl_2_ (purple), b) ATP/10 mM MgCl_2_, c) GTP/10 mM MgCl_2_, d) ADP/10 mM MgCl_2_/10 mM phosphate (Pi), and e) GDP/10 mM MgCl_2_/10 mM Pi. f) Time-dependent ATP (blue) and GTP (red) hydrolysis by TBCK. The formation of Pi was monitored by tracking absorbance at 360 nm over 4 minutes. The background signal from the negative controls without nucleotides has been subtracted from each reaction. Positive control is represented in black.

### 3.3 TBCK does not have ATPase or GTPase activities

Since some protein kinases also show intrinsic ATPase activity independent of substrate phosphorylation^23^, we tested whether TBCK can hydrolyze ATP *in vitro*. This activity can be measured by tracking phosphate production in real-time using an enzymatic coupling assay.^24^ Briefly, phosphate production was detected by UV-Vis through phosphorylation of 2-amino-6-mercapto-7-methylpurine riboside substrate, which is catalyzed by the purine nucleoside phosphorylase, resulting in ribose 1-phosphate and 2-amino-6-mercapto-7-methylpurine products. The real-time spectrophotometric shift was recorded in the presence or absence of 5 mM ATP, a concentration that is considered high under physiological conditions. A known reaction that produces inorganic phosphate (see Materials and Methods) was used as a positive control to confirm the function of the coupling reagents. A time-dependent increase in phosphate production was only observed in the positive control, but not in the TBCK-containing sample (**Figure 4f**). This indicates that the TBCK pseudokinase domain exhibits no detectable ATPase activity under these conditions, likely due to the lack of nucleotide/divalent cation binding or the catalytic motifs for hydrolysis. Similarly, no GTP hydrolysis was observed in the assay when ATP was replaced with GTP, indicating this is not specific to ATP.

## 4. Discussion

TBCK is a ‘dark’ protein kinase that plays a crucial role in vital cellular pathways, including mTOR signaling, autophagy, lysosomal function, mitochondrial fission, cell proliferation, and intracellular vesicle trafficking ^25^. Characterizing the biochemical and biophysical properties *in vitro* lays the foundation for understanding the TBCK-associated disease.

Through bioinformatics analysis, nearly 900 mutations have been reported to date, with more than 100 classified or predicted as pathogenic or likely pathogenic ^7,26^. Due to limited awareness and access to genomic testing and neuroimaging, the number of TBCKE cases is likely underreported ^4,27^. This is highlighted by two recent studies that retrospectively analyzed tissue from individuals who died in childhood decades ago ^28,29^. Emerging evidence further suggests that *TBCK* heterozygous carriers may present with neurological and cardiac abnormalities, reflecting the TBCK’s fundamental role in developmental processes ^30,31^. In line with these observations, animal models such as *TBCK*^+/−^ mice and *TBCK*^-/−^ knockdown zebrafish exhibit axonal neuropathy and defects in organ development, reinforcing TBCK’s fundamental importance in the neurodevelopmental processes ^30,31^.

Our bioinformatic studies suggest that the pseudokinase domain of TBCK is a class I pseudokinase, lacking the essential nucleotides and divalent cation-binding sites, as well as the motifs required for catalytic competence as a protein kinase. The AlphaFold4-predicted structure, although key regions in the pseudokinase cavity exhibit low confidence in the prediction score, does present a sizable cavity that could accommodate nucleotides. Our experimental characterizations show no ligand-induced stabilization at elevated temperatures. The activity assay also rules out the possibility that TBCK could hydrolyze ATP or GTP *in vitro*. These observations are consistent with the absence of canonical nucleotide-binding and catalytic motifs, supporting the hypothesis that TBCK is a class I pseudokinase.

As a multidomain pseudokinase, the biological function of TBCK is likely governed by contributions from its auxiliary domains. Intriguingly, a large number of disease-associated missense mutations are localized within the TBC domain rather than the pseudokinase domain, highlighting its potential functional importance ^7^. The TBC domain of TBCK retains conserved arginine and glutamine residues characteristic of GTPase-activating proteins (GAPs), suggesting that TBCK may function as a GAP toward small GTPases ^7,32^. In contrast, the rhodanese-like domain is predicted to be catalytically inactive, as it lacks the conserved CX_4-5_R active-site motif found in enzymatically active rhodanese family members, based on comparisons with structurally related proteins ^33^. The precise roles of the TBC and rhodanese-like domains remain to be defined. Nonetheless, our work establishes the first biophysical and biochemical framework for TBCK.

## 5. Conclusions

This work presents the *in vitro* characterization of the human TBCK, establishing a foundation for elucidating its molecular function in TBCKE. Our data support the classification of TBCK as a class I pseudokinase, lacking nucleotide binding and hydrolytic activity. These findings provide a framework for future mechanistic studies of TBCK biochemical function and may ultimately contribute to improved diagnosis and the development of targeted therapeutic strategies for individuals with TBCK-related disorders.

## Author Contributions

SM: conceptualization, investigation, *in silico* and *in vitro* experiments, formal analysis, visualization, writing initial draft, and editing. ATI: mass spectrometry and editing. LC: limited proteolysis experiments and writing the initial draft. WZ: conceptualization, investigation, editing, formal analysis, supervision, and critical review of the manuscript.

## Acknowledgement

This work was supported by the National Institutes of Health (R35GM160337 to WZ and 1S10OD020062-01 to ATI), the Florida State University (FSU) Start-up Fund, and the FSU Institute for Pediatric Rare Disease Award. The authors thank Drs. Peter Randolph, Gwimoon Seo, Brian K. Washburn, and John M. Schulze at the FSU Institute of Molecular Biophysics Protein Biophysics Facility, FSU Institute of Molecular Biophysics Protein Expression Facility, FSU Department of Biological Science Molecular Cloning Facility, and University of California Davis Molecular Structure Facility, respectively, for training and technical support.

